# Characterisation of the role and regulation of *Ultrabithorax* in sculpting fine-scale leg morphology

**DOI:** 10.1101/2020.06.17.152918

**Authors:** Alexandra D. Buffry, Sebastian Kittelmann, Alistair P. McGregor

## Abstract

Hox genes are expressed during embryogenesis and determine the regional identity of animal bodies along the antero-posterior axis. However, they also function post-embryonically to sculpt fine-scale morphology. To better understand how Hox genes are integrated into post-embryonic gene regulatory networks, we further analysed the role and regulation of *Ultrabithorax* (*Ubx*) during leg development in *Drosophila melanogaster. Ubx* regulates several aspects of bristle and trichome patterning on the femurs of the second (T2) and third (T3) leg pairs. We found that repression of trichomes in the proximal posterior region of the T2 femur by Ubx is likely mediated by activation of the expression of *microRNA-92a* by this Hox protein. Furthermore, we identified a novel enhancer of *Ubx* that recapitulates the temporal and regional activity of this gene in T2 and T3 legs. We then used transcription factor binding motif analysis in regions of accessible chromatin in T2 leg cells to predict and functionally test transcription factors that may regulate the *Ubx* leg enhancer. We also tested the role of the Ubx co-factors Homothorax (Hth) and Extradenticle (Exd) in T2 and T3 femurs. We found several transcription factors that may act upstream or in concert with Ubx to modulate trichome patterning along the proximo-distal axis of developing femurs and that the repression of trichomes also requires Hth and Exd. Taken together our results provide insights into how *Ubx* is integrated into a postembryonic gene regulatory network to determine fine-scale leg morphology.

## Introduction

The Hox genes encode an important and conserved family of transcription factors (TFs) that are expressed during embryogenesis to determine the identity of body regions along the anteroposterior axis of animals [1–4]. Hox genes also play more subtle but important post-embryonic roles in regulating cell identify to sculpt the fine-scale morphology of structures and organs, and consequently have been likened to ‘micromanagers’ [5–8]. Several such post-embryonic roles of Hox genes have been identified in *Drosophila*; for example, the specification of certain subtypes of cells in the central nervous system [9,10], the regulation of the development of larval oenocytes by *abdominal-A* (*abd-A*) [11], and the integration of regulatory information to specify differences in prothoracic (T1) leg bristle patterning among leg segments and between sexes by *Sex-combs reduced* (*Scr*) [12].

*Ultrabithorax* (*Ubx*) specifies the identity of thoracic and abdominal segments during *Drosophila* embryogenesis [13–16]. Classically, this Hox gene represses wing identity and promotes haltere formation on the third thoracic (T3) segment through the direct regulation of potentially hundreds of genes [15–19]. *Ubx* also distinguishes the size and morphology of halteres at a more fine-scale level, in part through the autoregulation of differences in the expression levels between proximal and distal cells [20,21]. In these appendages *Ubx* also influences chromatin accessibility through cell type-specific interaction with co-factors and can thereby act as a repressor as well as an activator [22]. During mesothoracic (T2) and T3 leg development, Ubx is expressed along the proximo-distal axis of pupal femurs, with the highest concentration in proximal posterior and dorsal-anterior cells [23,24]. This expression of Ubx regulates the patterning of trichomes and bristles on the T2 and T3 femurs in a concentration dependent manner [23–31]. Therefore, in addition to determining segmental identity, *Ubx* subsequently contributes to sculpting the finer-scale morphology of several appendages.

Despite these insights into Hox gene function, we still do not fully understand how they are integrated into post-embryonic gene regulatory networks (GRNs). One approach to address this is to study the regulation of Hox genes by identifying the enhancers that are responsible for their post-embryonic expression. Indeed, several enhancers and other cis-regulatory elements of *Ubx* have already been identified and we are beginning to understand how they integrate information to precisely regulate the differential expression of this Hox gene to control fine-scale morphology [20,32–41] However, it is clear that not all *Ubx* enhancers have been identified and we still have much to learn about the complex regulation of this crucial gene [20,23,36].

Enhancers can be challenging to identify because currently there is no consensus of what genomic features mark these regions [42,43]. Furthermore, although the regulatory genome can now more readily be studied with new tools such as ATAC-seq, C technologies and CRISPR/Cas9, we still do not fully understand the regulatory logic underlying enhancer function [42–45]. Given their importance in development, disease and evolution, it is crucial that we continue to identify and study individual enhancers in detail, to better our general understanding of cis-regulatory regions and GRNs.

The development and patterning of trichomes among *Drosophila* species has proven an excellent model to study GRNs and their evolution [46–48]. Trichomes are short, non-sensory actin protrusions that are found on insect bodies throughout all stages of life [46]. They are thought to be involved in processes such as aerodynamics, thermal regulation and larval locomotion [49,50]. The larval cuticle of *Drosophila* displays a distinct pattern of trichomes and the underlying GRN is understood in great detail [51–53]. In brief, the gene *shavenbaby* (*svb*) appears to integrate information from upstream factors, including Ubx, and directs expression of downstream effector genes that determine the formation of the trichomes themselves [51–55]. Moreover, the convergent evolution of larval trichome patterns in different *Drosophila* lineages is caused by changes in enhancers of *svb* [47,54–60].

The T2 legs of *D. melanogaster* display a trichome pattern with a patch of cuticle on the proximal posterior of the femur that is free from trichomes, known as the “naked valley” (NV) [24,61] (Fig. 1A,C). We previously studied the GRN underlying leg trichome patterning and found that it differs in topology with respect to the larval trichome GRN [62]. In particular, in the developing T2 legs, the Svb-dependent activation of trichomes is blocked by microRNA-92a (miR-92a)-mediated repression of Svb target genes to generate the NV [61,62]. Furthermore, in contrast to its activation of the larval trichomes, Ubx represses leg trichomes perhaps via miR-92a [24,54,61].

**Figure 1.**
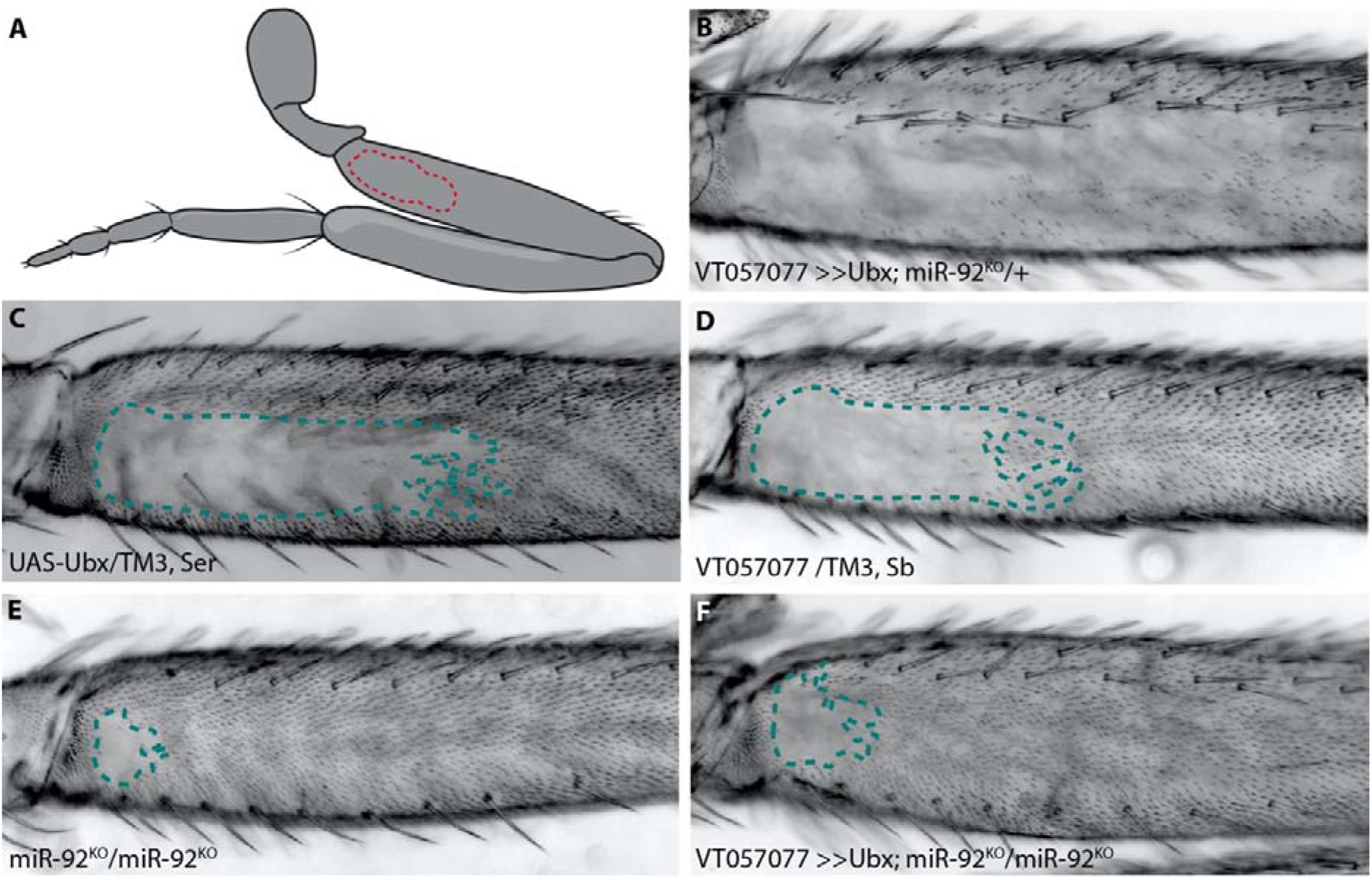
*Ubx* requires *miR-92a* to repress trichomes. The naked valley is a region of trichome-free cuticle on the posterior side of the proximal femur (outlined by dotted lines) (A). Over-expression of *Ubx* inhibits trichome formation on most of the T2 femur (B) while the *UAS-Ubx* line (C) and the GAL4 driver line VT057077 (D) both have large naked valleys. Naked cuticle is almost absent in *miR-92* loss of function T2 legs (E). Trichome inhibition by *Ubx* over-expression is restricted to the most proximal region of the femur in *miR-92*^KO^ (F).

The size of the NV varies within and between species and these differences are associated with changes in the expression of *miR-92a* [61] and *Ubx* [24], respectively. Ubx is expressed in *D. melanogaster* T2 legs in the region of the NV, but not in the T2 legs of *D. virilis*, which has no NV [24]. Moreover, it has been shown that *Ubx* contributes to differences in NV size between *D. melanogaster* and *D. simulans* [24]. It was postulated that the evolution of *Ubx* expression in T2 legs is attributable to the presence of a T2 leg-specific enhancer of *Ubx* [23]. However, no cis-regulatory sequences that could drive expression of *Ubx* in T2 were identified.

Here we further characterise how Ubx is wired into the GRN for leg trichome patterning. We show that repression of trichomes by Ubx is likely dependent on *miR-92a* activation by this Hox gene. We also identified a novel enhancer of *Ubx* that drives expression along the proximo-distal axis of T2 and T3 femurs during trichome patterning. Functional analysis of TFs predicted to bind to this *Ubx* leg enhancer revealed that several activate or repress leg trichomes and that repression of trichomes by Ubx is dependent on the co-factors Extradenticle (Exd) and Homothorax (Hth). Taken together our results provide new insights into the role and regulation of *Ubx* during post-embryonic development and in sculpting fine-scale adult morphology.

## Materials and Methods

### Fly stocks and genetics

All stocks used were kept on standard yeast extract-sucrose medium at 25°C. Reporter lines VT42732, VT42733 and VT42734 were obtained from the Vienna *Drosophila* Resource Centre (VDRC). Reporter lines GMR31F12, GMR32BO3 and GMR31E11 were obtained from the FlyLight enhancer collection [63]. To test the activity of all enhancer lines, they were crossed to a UAS-stingerGFP and/or UAS-shavenoid(sha)ΔUTR. To test the interaction between *Ubx* and *miR-92a*, we crossed UAS-Ubx flies to a pan-epidermal GAL4 driver (VT057077; VDRC) in a *miR-92* loss-of-function background [64]. To test putative transcription factors that bind to VT42733, UAS-RNAi lines for candidate transcription factors were crossed to VT42733. A list of all stocks used can be found in S1 File.

### Cloning

Fragments UbxP1, e33.A, e33.B and e33.C were PCR-amplified from genomic DNA (*D. melanogaster*, Oregon R). UbxP1 was cloned directly into the S3aG expression vector (a gift from Thomas Williams, Addgene plasmid #31171). Fragments, e33.A, e33.B and e33.C, were initially inserted into the TOPO/D vector (Invitrogen). LR gateway cloning was then used to subclone the fragments into the pBPGUw plasmid upstream of GAL4 (a gift from Gerald Rubin, Addgene plasmid #17575). The resulting constructs were inserted into landing site 86Fb using phiC31 mediated germline transformation by either BestGene Inc or the Cambridge injection facility. Genomic coordinates of fragments and primer sequences can be found in S2 File.

### GFP and NV analysis

To assay expression in pupae, white prepupae from GAL4 lines crossed to UAS-stingerGFP were collected and aged to between 20 and 28 hours after puparium formation (hAPF), the window when T2 trichome patterning is regulated by *Ubx* [24]. UbxP1 flies were analysed without crossing to UAS-GFP as they were constructed in such a way to allow direct GFP expression. GFP expressing whole pupae were imaged on a Zeiss Axiozoom stereoscope. For the dissection of pupal legs, the pupal case was removed and the pupae were covered in 4% formaldehyde for 10 mins, a small dissection needle was then used to create small openings in the head and abdomen. Fresh 4% formaldehyde was flushed over the pupae and left for another 10 minutes. T2 and T3 pupal legs were then dissected with tungsten needles and mounted in 80% glycerol. Mounted legs were immediately imaged on a Zeiss 800 confocal. For the analysis of expression patterns in T2 and T3 leg discs, 3^rd^ instar larvae were dissected and reporter expression visualised with anti-GFP (Thermofisher) (1:600) and goat anti-chicken 488 (1:600) according to standard protocols. Discs were also stained with DAPI and mounted in 80% glycerol and imaged on a Zeiss 800 confocal. For the analysis of trichome patterns, T2 and T3 legs were dissected from adults and mounted in Hoyer’s medium/lactic acid (1:1) and imaged under a Zeiss Axioplan microscope with a ProgRes MF cool camera (Jenaoptik). The size of the NV was measured (n = at least 10) using Fiji software [65] and statistical analysis was performed in R-Studio version 1.2.1335 [66]. We expect the NV of the progeny to be an intermediate size between the two parental lines, if this was not the case further statistics were carried out. Data was checked for normality using Shapiro-Wilk, followed by either an ANOVA or Kruskal-Wallis test. To check significance between each group either Tukey’s post-hoc test or Dunn’s test was performed, p-values were adjusted using Bonferroni correction to avoid multiple testing errors. For SEM imaging, legs were dissected in adult flies and stored in fresh 100% ethanol. Legs were then critically point dried using automatic mode of a Tousimis 931.GL Critical Point Dryer, mounted on SEM stubs with carbon tabs, sputter-coated with a 15 nm thick gold coat and imaged in a Hitachi S-3400N at 5kV with secondary electrons.

### Identification and functional testing of candidate TFs

To identify potential TFs that bind to the *Ubx* leg enhancer, the JASPAR TF database was utilised [67] with a relative profile threshold of 85% similarity. The resulting factors were compared to the RNA-seq data for T2 legs (GEO accession number GSE113240) [62], and genes encoding TFs with an expression level of over 1 fragment per kb per million (FKPM) were scored as expressed. To further filter TFs, only those with predicted binding sites in regions of accessible chromatin, from T2 leg ATAC-seq profiling data (S3 File) (GEO accession number GSE113240) [62] were selected. To assay whether the identified TFs have any role in trichome development on the T2 and T3 legs, RNAi lines for selected genes were crossed to the Ubx VT33 enhancer and the resulting trichome pattern was measured and compared to parental control lines (S4 File).

## Results

### Ubx repression of T2 leg trichomes requires miR-92a

In addition to its well characterised role in T3 leg development, it was previously found that *Ubx* represses the formation of trichomes on T2 femurs in a dose sensitive manner from proximal to distal [24]. We corroborated this finding by over expressing *Ubx* in T2 legs, which resulted in loss of all proximal and most of the distal trichomes on posterior T2 femurs, including those dorsal and ventral of the NV (Fig. 1A-D). As we showed previously, over-expression of *miR-92a* also represses T2 trichomes and, reciprocally, loss of this microRNA results in a very small NV [61,62] (Fig. 1E). This suggests that Ubx acts upstream of *mir-92a* to inhibit trichome formation. In order to test this, we over expressed *Ubx* in flies homozygous for a loss of function of *mir-92a* (and its paralogue *miR-92b*) [64]. We found that *Ubx* is unable to repress trichomes in the absence of *mir-92a* (Fig. 1F).

We also tested the effects of *Ubx* over-expression and miR-92 loss of function on T3 leg trichomes (S1 Fig). Without miR-92a, trichomes develop in normally naked regions of the posterior T3 femur, albeit in a patchy pattern (S1 Fig). This is also the case when *Ubx* is overexpressed in the absence of mir-92, indicating again that Ubx requires *miR-92a* to repress trichome development on the posterior of T3 femurs. Note that Ubx over-expression never interferes with the formation of anterior trichomes on T2 or T3 femurs (S1 Fig). Taken together, our findings suggest that *Ubx* represses trichomes on posterior femurs by directly or indirectly activating *miR-92a* expression, which in turn inhibits the expression of Svb target genes including *shavenoid* (*sha*) [61,62,68]. To better understand how *Ubx* is integrated into the leg trichome GRN, we next attempted to identify cis-regulatory elements that regulate expression of this Hox gene in T2 and T3 legs.

### Several regions of the Ubx locus with open chromatin drive expression in Drosophila pupal legs

To identify the previously predicted leg *Ubx* enhancer, Davis et al. [23] assayed available regulatory mutations of the *Ubx* locus and generated new deficiencies. This allowed them to rule out approximately 100 kb in and around the *Ubx* locus as containing the T2 leg enhancer. They then assayed a further 30 kb using reporter constructs (Fig. 2A). In total they investigated over 95% of the *Ubx* locus but were unable to identify a region with T2 leg specific activity.

**Figure 2.**
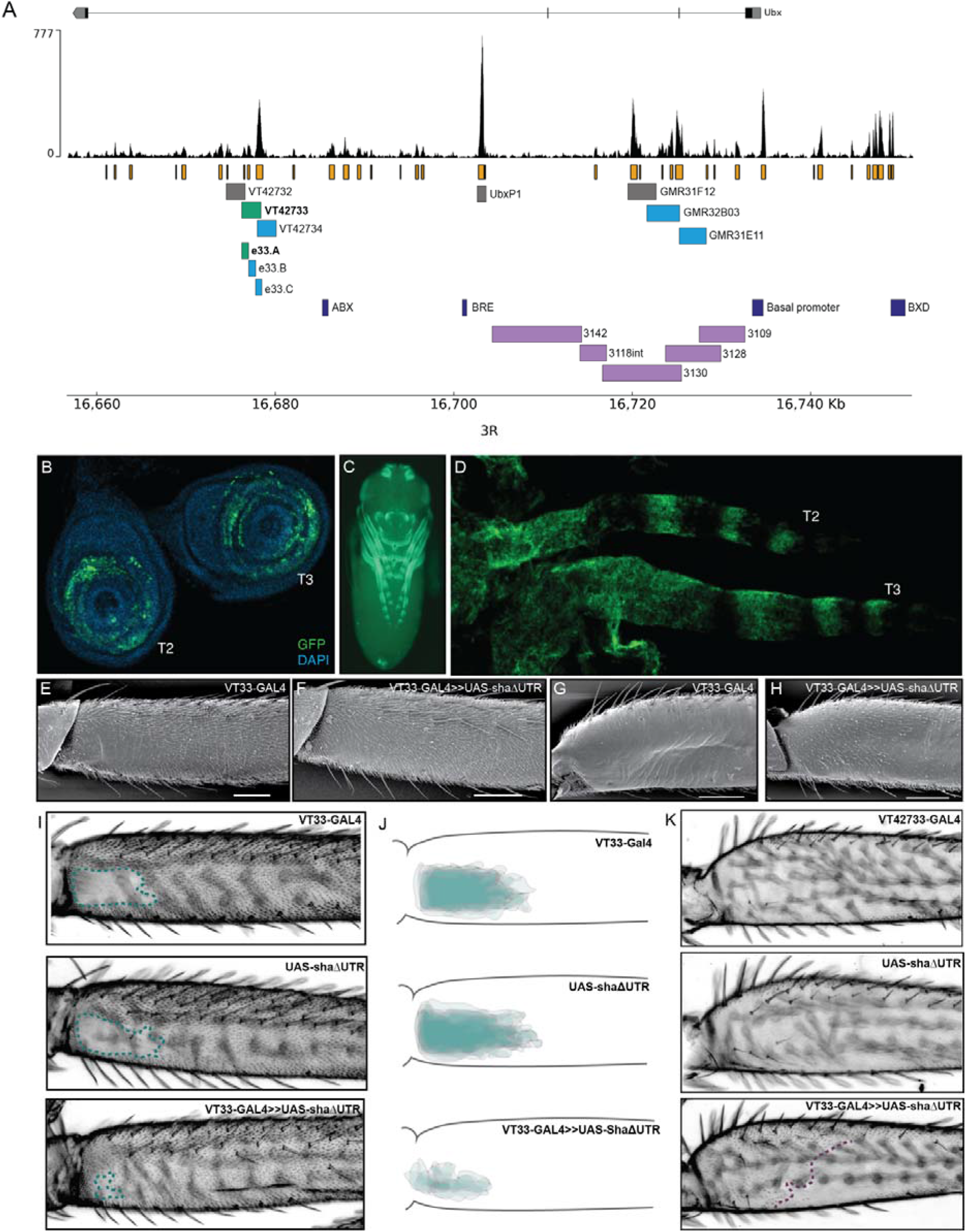
Testing regions of accessible chromatin at the *Ubx* locus for enhancer activity. (A) The *Ubx* locus in *Drosophila melanogaster* on chromosome arm 3R with the ATAC-seq profile for *Ubx* in pupal legs below. In orange are peaks of open chromatin and underneath are the locations of tested or known regulatory elements. In green are the lines tested in this study that affect trichome patterning on T2 and T3 legs. In blue are the lines which express GFP in pupal legs but do not have a functional effect on trichome patterning. In grey are the lines which do not drive expression in pupal legs. In dark blue are a subset of the characterised regulatory elements of *Ubx*. For a complete list of all characterised elements according to RedFly see S2 File. In purple are the reporter constructs that were tested by Davis et al., [23] that did not contain the leg specific enhancer of *Ubx*. (B) UAS-GFP expression driven by VT42733-GAL4 (referred to as VT33-GAL4) in T2 and T3 leg imaginal discs from 3rd instar larvae is seen in the femur ring of the disc but also other leg segments. (C) Expression of VT33 in whole pupae at 24 hAPF. Expression is quite promiscuous and can be seen in the pupal legs, antennae, mouthparts, eyes, and genitalia. (D) In T2 and T3 pupal legs from flies at 24 hAPF, the expression driven by VT33 is observed in the developing femur and also in a striped pattern more distally in the pupal leg. (E) Scanning electron micrograph (SEM) showing the NV of a wild-type T2 proximal femur. (F) SEM of a T2 femur when the VT33 enhancer is crossed to the trichome activating line, UAS-shaΔUTR. Most of the naked cuticle is now filled with trichomes. (G) SEM of a wild-type T3 femur shows that the posterior region of the leg is largely free of trichomes. (H) When the VT33 enhancer is crossed to UAS-shaΔUTR, there is ectopic development of trichomes on the proximal part of the T3 femur. (I) Light-microscope images of adult legs from progeny of the VT33 enhancer crossed to UAS-shaΔUTR. The parental controls are shown on top and the progeny of the cross beneath. In each case the naked valley is outlined with a green dashed line. (J) Visual representation of VT33 GAL4 crossed to UAS-shaΔUTR and controls (n=10). There is a dramatic decrease in the size of the NV in all individuals. (K) Images of the T3 femur when VT33 is crossed to the trichome activating line versus the parental control lines.

To follow up the work of Davis et al. [23], we used ATAC-seq data that we generated previously to identify regions of accessible chromatin in T2 leg cells in the developmental window when the trichome pattern is determined [62] (Fig. 2A). We found that the *Ubx* locus contains several regions of accessible chromatin in T2 pupal cells corresponding to known enhancers or promoters as well as putative new cis-regulatory elements (Fig. 2A; S2 File).

We then took advantage of existing reporter lines [63,69], to assay regions of open chromatin in the introns of *Ubx* for enhancer activity, and specifically to test if any could drive expression in developing T2 legs (Fig. 2A). We selected three lines from the GMR database that overlap with lines tested by Davis et al. [23], but did not show enhancer activity in T2 legs (Fig. 2A). We also assayed three lines from the VT-GAL4 database (VDRC) corresponding to several peaks of open chromatin but not overlapping with any known regulatory elements of *Ubx* [RedFly: 70] (S2 File). Finally, we tested the UbxP1 peak of accessible chromatin, which corresponds to a previously characterised variably occupied CTCF site [36]. This region was not covered by Davis et al., [23] and therefore was not previously tested for enhancer activity in legs. Four of the seven regions tested were able to drive reporter gene expression in developing legs at 24 hAPF, although all of them appeared to be quite promiscuous and had activity in other pupal tissues (Fig. 2A,C and Fig. S2).

We next tested whether regions VT42733, VT427734, GMR32B03 and GMR31E11 (Fig. 2A), which drive GFP expression in pupal legs, could also influence the trichome pattern on the femurs of T2 and T3 legs, which would further indicate that they are active in leg epidermal cells at the time of trichome patterning. To do this we crossed the driver lines to UAS-shaΔUTR, which overrides trichome repression by miR-92a and leads to the formation of trichomes on normally naked cuticle [61]. Therefore, in this assay, enhancer regions that are active in posterior femurs at the correct time will generate trichomes where there is normally naked cuticle. Only one of the reporter lines identified, VT42733, was able to induce the formation of trichomes in the NV, resulting in a striking decrease in the size of the patch of naked cuticle (Fig. 2A,E,F,I,J). Importantly, we noticed that while VT42733 greatly reduces the size of the NV in this assay, a small patch of naked cuticle remains proximally on the ventral side of the T2 posterior femur (Fig. 2F,I,J), which is consistent with Ubx-independent repression of trichomes in these cells [23]. We also observed that VT42733 was able to induce the formation of trichomes proximally on the posterior and dorsal-anterior of T3 femurs, suggesting that this enhancer also contributes to T3 femur patterning (Fig. 2G,H,K). The proximal dorsal-anterior activity of VT42733 in T3 femurs overlaps with the activity of *abx* [23]. We observed that the activity of VT42733 in the T3 femur is proximally restricted and does not extend as distally as where Ubx is known to repress trichomes, which is consistent with previous data showing that *pbx* and potentially *bx* also regulate expression of this Hox gene in the posterior of T3 femurs [23].

We examined the expression driven by VT42733 in more detail in leg imaginal discs and in pupal legs (Fig. 2B,D). In 3^rd^ instar leg discs, GFP expression driven by this enhancer can clearly be seen in rings which will develop into the future T2 and T3 femurs (Fig. 2B). Similar reporter expression can be seen in the pupal T2 and T3 femurs, as well as more distal segments (Fig. 2D), which is consistent with the fact that this region can promote trichome formation on T2 and T3 femurs when combined with UAS-shaΔUTR (Fig. 2E-K).

Taken together these results evidence that VT42733 represents a novel *Ubx* leg enhancer, which regulates expression of this Hox gene in the NV region of T2 femurs as well as proximally in T3 femurs.

### Delineation of the Ubx leg enhancer

VT42733 drives expression in T2 and T3 legs consistent with Ubx activity, but this enhancer is also active in other pupal tissues (Fig. 2C). To further delineate the *Ubx* leg enhancer region, VT42733 was broken down into three partially overlapping fragments of around 700 bp: e33.A, e33.B, and e33.C (Fig. 2A). All three lines were able to drive reporter expression in developing pupae (Fig. 3 and S2 Fig): e33A drives expression in leg discs, pupal legs, antennae and developing eyes (Fig. 3A,B), and e33.B drives a more restricted expression pattern limited to a small patch in the pupal legs and in the head (S2 Fig). While e33.C also drives expression in the legs, its activity is predominantly in the head and thorax as well as a stripe-like pattern on the ventral side of the abdomen, which was not seen with any of the other driver lines tested (S2 Fig).

**Figure 3.**
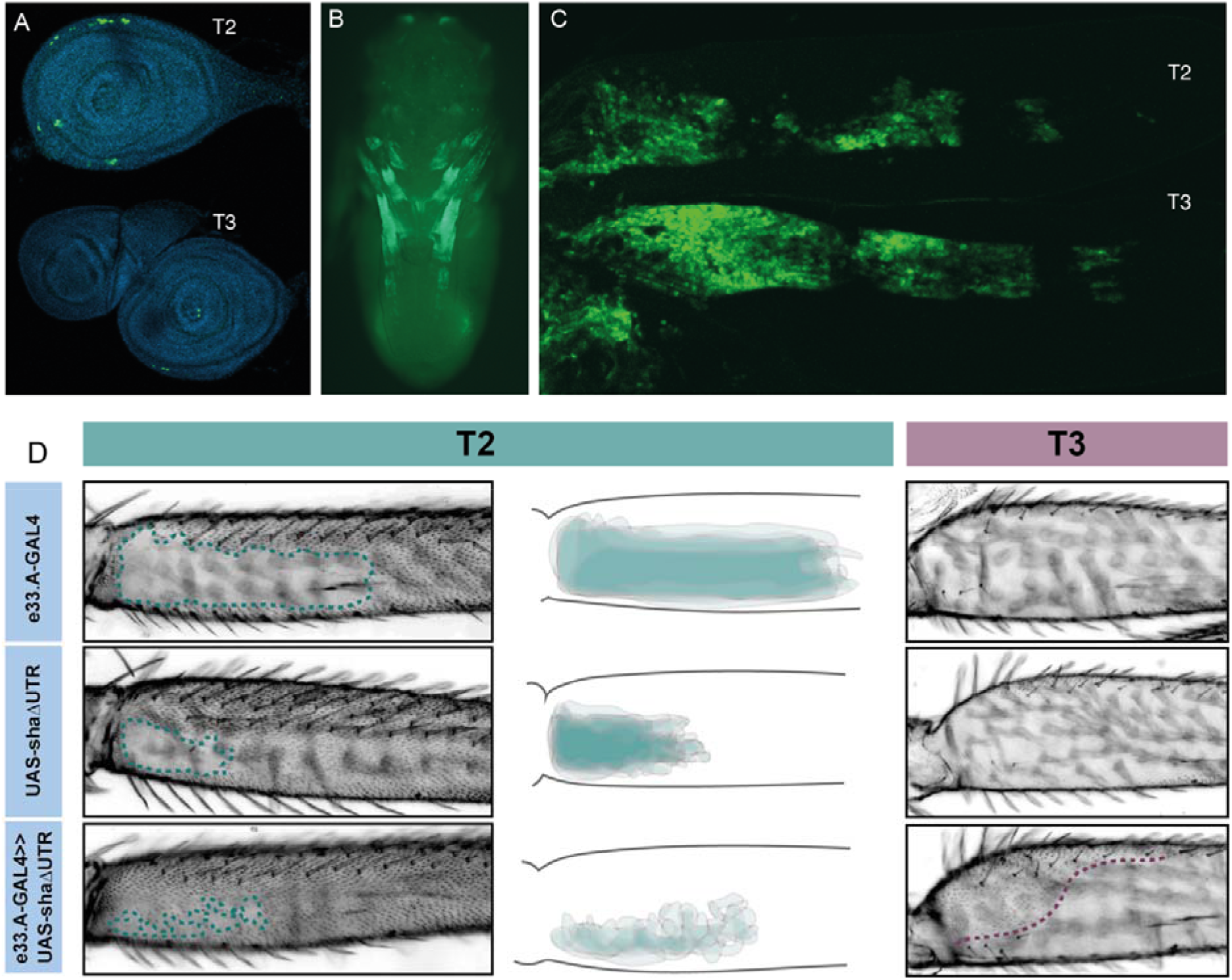
Characterisation of e33.A enhancer activity. (A) Expression driven by e33.A in T2 and T3 leg imaginal discs from 3^rd^ instar larvae is restricted in comparison to the whole VT33 enhancer, and expression. There is also reduced expression in T3 compared to T2. (B) In whole pupae at 24 hAPF e33.A drives mainly in the developing legs. (C) In T2 and T3 pupal legs at 24 hAPF, there is clear expression of e33.A in the femur. (D) The result of crossing e33.A to the trichome activating line, UAS-shaΔUTR. On T2 legs (labelled in green), there is patchy ectopic trichome development in the NV when compared with parental controls. There is also ectopic trichome development in the proximal part of the T3 femur (outlined in purple), although this does not extend as far ventrally as with the VT33 enhancer.

To further test the functionality of these smaller fragments, they were combined with UAS-shaΔUTR. We observed that e33.A was able to drive trichomes in the T2 NV albeit in a patchy and irregular pattern compared to VT42733 (Figs 2 and 3). e33.A also drove trichomes proximally in the dorsal-anterior and posterior of T3 femurs although this activity did not extend as far distally as with VT42733 (Figs 2 and 3). e33.B and e33.C did not have any detectable activity in this assay (S2 Fig). This suggests at least part of the enhancer activity of VT42733 in developing T2 and T3 femurs is determined by TF binding sites (TFBS) in e33.A.

### Analysis of transcription factors that may bind to the Ubx leg enhancer

To further characterise the *Ubx* leg enhancer, we carried out motif analysis to identify TFs that may bind to this region. To focus on binding sites for TFs that are expressed at the time of trichome development, we cross-referenced previously generated RNA-seq data for T2 legs [62] with the JASPAR database [67] (with the caveat that the JASPAR database does not contain an exhaustive list of all *Drosophila* TFs). Using a threshold of 85% similarity and focussing only on TFs expressed above a 1 FPKM threshold in T2 legs, 62 TFs were found to have predicted binding sites in the VT42733 region. We then further filtered the TFs using T2 pupal leg ATAC-seq data [62] to shortlist TFs with predicted binding sites located only in the accessible chromatin of region VT42733. This resulted in a total of 55 TFs (35 with predicted binding sites in e33.A) that are expressed in pupal T2 legs and predicted to bind to accessible regions in the VT42733 enhancer (S3 File).

We then tested the role of 25 of these TF candidates, as well as the known Ubx co-factor *homothorax* (*hth*) in T2 and T3 femur patterning by knocking-down their expression using RNAi (S3,S4 Files). To do this we used VT33-GAL4 since it expresses in the NV cells in the correct window of pupal leg development. We found that knockdown of 8/26 genes affected the trichome pattern on the posterior of T2 femurs when normalised for femur length: *arrowhead* (*awh*), *C15, Distal-less* (*Dll*), *extradenticle* (*exd*), *hth, mirror* (*mir*) *NK7.1* and *ventral-veins lacking* (*vvl*) (Figs 4–6, S3, S4 Figs, and S4 File).

**Figure 4.**
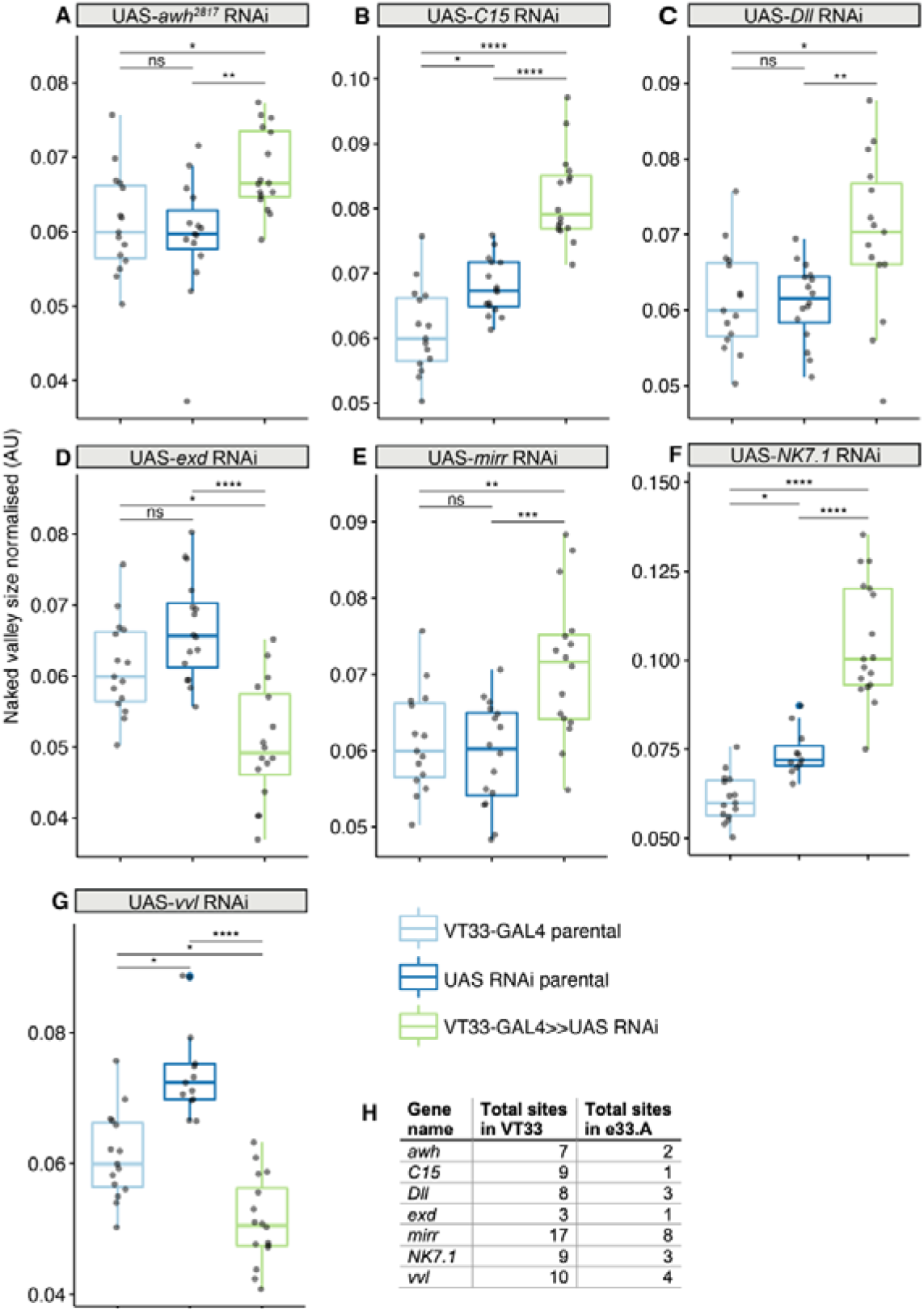
Transcription factors predicted to bind the *Ubx* leg enhancer that significantly affect T2 trichome patterning. Knockdown of *awh* (A), *C15* (B), *Dll* (C), *mirr* (E) and *NK7.1* (F) results in a significant increase in the size of the NV, while *exd* (D) and *vvl* (G) knockdown makes the NV smaller. In each case the progeny from the cross between the UAS-RNAi and the VT33-GAL4 (green boxes) were compared to the parental strains (blue boxes). Significance levels are shown above the pairs, p-values (p > 0.05 NS, P ≤ 0.05 *, P ≤ 0.01 **, P 0.001 ***, P ≤ 0.0001 ****) were calculated with an ANOVA or Kruskal-Wallis depending on normality. (H) Summary of the number of binding sites found by JASPAR for the seven TFs in the whole of the VT33 enhancer and how many of those sites are found in e33.A.

**Figure 5.**
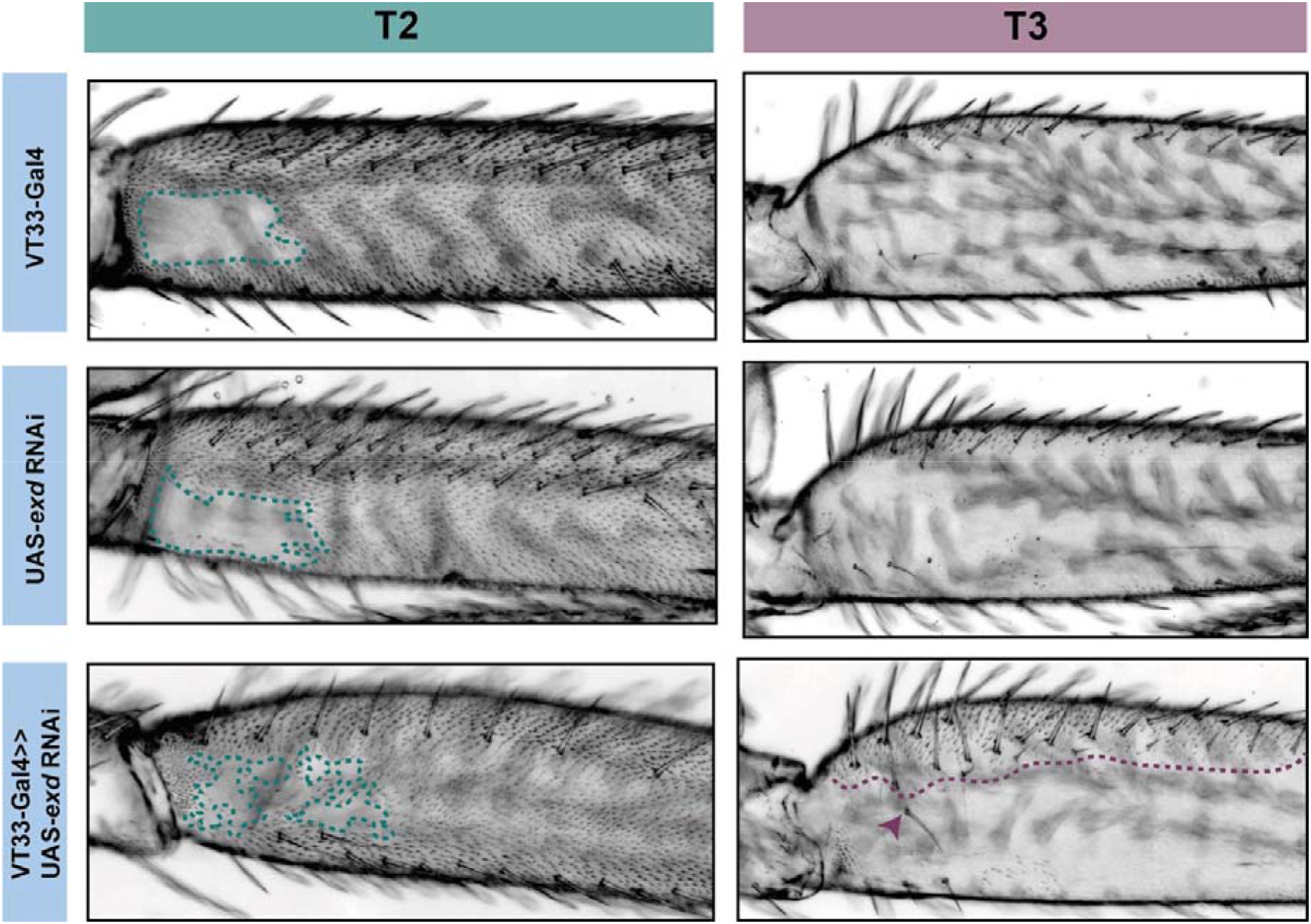
RNAi knockdown of *exd* results in ectopic trichome growth. *exd*-RNAi results in patchy ectopic trichomes in the NV of T2 femurs (green). On T3 femurs (purple), *exd*-RNAi also causes the development of ectopic trichomes, which extend about one third from the proximal dorsal towards the ventral (purple dotted line). There is also an additional row of bristles as well as ectopic bristle growth (arrowhead).

**Figure 6.**
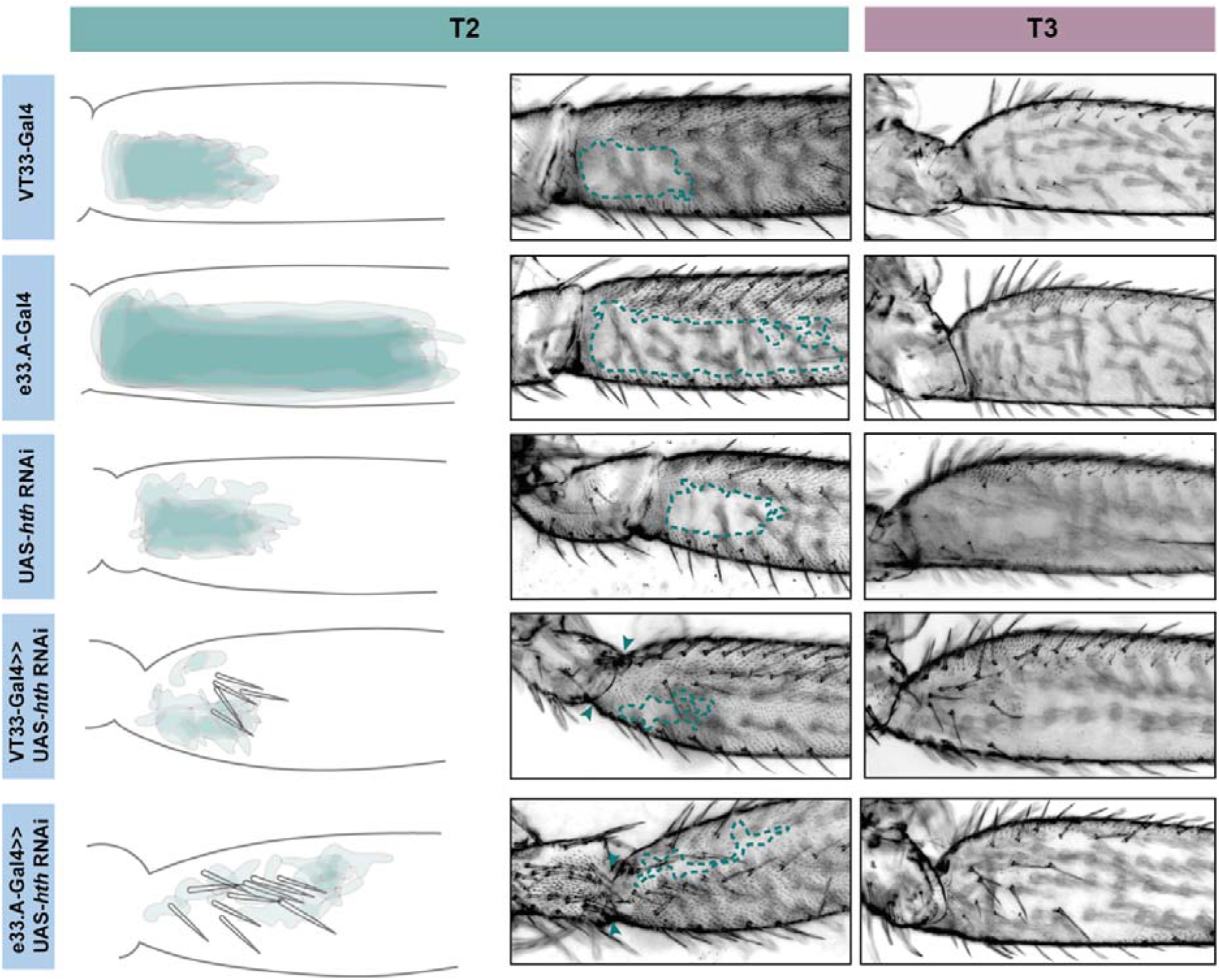
Knockdown *hth* affects leg morphology and patterning. In T2 (green) knockdown of *hth* causes the ectopic development of trichomes in the NV as well as sensory bristles in the NV. There is also a change in the shape of the coxa and femur suggesting a transition to a more T1 like phenotype. This effect is more severe when the knockdown is driven with e33.A-GAL4; there is also the development of trichomes and bristles on the coxa. On T3 (purple) knockdown of *hth* similarly results in ectopic sensory bristles and trichomes that are restricted to the proximal of the femur.

Knockdown of *exd* and *hth* resulted in ectopic trichomes at the proximal posterior of T2 femurs (Figs 5 and 6). *hth* knockdown had a stronger effect than *exd* on T2 morphology, with ectopic sensory bristles and transformation of the shape of the coxa to a more T1 like appearance (Fig. 6). Surprisingly these effects were even more pronounced when using e33.A as a driver (Fig. 6). On T3, knockdown of *exd* resulted in trichome formation in the dorsal-anterior of the femur and additional bristles along the A-P boundary (Fig. 5). Knockdown of *hth* again had a stronger effect on T3 with additional trichomes and bristles on the dorsal-anterior and proximally on the posterior femur (Fig. 6). These results are consistent with loss of *Ubx* function in T2 and T3 femurs [23] and suggest that Hth and Exd promote Ubx activity in T2 and T3 perhaps by acting as co-factors for this Hox gene in this context.

*vvl* was the only other TF tested that resulted in ectopic trichome formation in the proximal posterior of T2 femurs when knocked down (Fig. 4 and S3, S4 Figs). RNAi against *vvl* also produced ectopic trichomes in the dorsal-anterior of T3 femurs (S5 Fig). This indicates that Vvl supresses trichome formation in T2 and T3 femurs perhaps by promoting Ubx expression or acting down stream of this Hox gene.

We found that RNAi knockdown of *awh, C15, Dll, mir* and *NK7.1* resulted in a distal expansion of the NV on T2 femurs (Fig. 4, S3, S4 Fig, and S4 File), but had no effect on T3 femur patterning (Fig. S6). This suggests that these TFs may repress Ubx activity in the distal of the posterior T2 femur or promote trichome formation in this region.

## Discussion

### Identification of a *Ubx* leg enhancer

We have found that Ubx represses trichomes on the femurs of T2 and T3 legs via *mir-92* (Fig. 7). We then sought to determine how this Hox gene is regulated in these appendages. We searched for a *Ubx* leg enhancer guided by regions of accessible chromatin in this tissue, identified using ATAC-seq. Four of the seven lines from existing reporter construct collections tested were able to drive detectable GFP expression in the developing pupal legs, but none of the lines drove expression specific to just the T2 and T3 femurs. However, using functional testing, we found that the 2.2 kb region VT42733 has enhancer activity in the proximal posterior of T2 and T3 femurs and proximal dorsal-anterior of T3 femurs during the correct developmental time point and consistent with Ubx functions in these legs (Fig. 2E-K). Analysis of sub-fragments of VT42733 showed that a 700 bp region, e33.A, is also active in T2 and T3 femurs cells, but this activity is weaker than the full VT42733 sequence (Fig. 3D) and has fewer predicted TF binding sites (S3 File). However, any additional binding sites in the region of VT42733 that does not overlap with e33.A appear insufficient to drive expression on their own, since e33.B and e33.C had no detectable functional activity. Taken together, these results indicate that the *Ubx* leg enhancer is located in region VT42733 with some binding sites concentrated in region e33.A (S3 File). Importantly, while VT42733 and e33.A are able to drive expression in the proximal femur, they are inactive in the ventral part of the posterior T2 and T3 femurs (Fig. 2F, I and Fig. 3D). This was particularly evident for VT42733 when combined with UAS-shaΔUTR, which resulted in the entire posterior T2 femur being covered in trichomes apart from a small ventral region (Fig. 2F, I). This is consistent with previous studies showing that while Ubx represses trichomes on the posterior T2 femurs, it is inactive in these ventral cells, and even in the absence of *Ubx*, this region of the cuticle fails to differentiate trichomes [23,28]. Indeed, this region also stays trichome-free in a miR-92 loss of function line (Fig. 2K, H), indicating that repression of trichomes in these cells is independent of *Ubx* and *miR-92a*. The expression driven by VT42733 is also consistent with Ubx activity in T3 femurs: repression of trichomes proximally on the posterior and on the proximal dorsal-anterior region [23] (Fig. 7). This suggests that the enhancer we have identified does indeed recapitulate the expression and activity of *Ubx* in T2 and T3 femurs. Interestingly, FAIRE-seq to assay the open chromatin in developing halteres and wings revealed that while the *abx* region is accessible there was no distinctive peak in the region of the new leg enhancer we have discovered here [20,71]. This suggests that while the enhancer we have identified is accessible and active in legs it is not used in the developing halteres.

**Figure 7.**
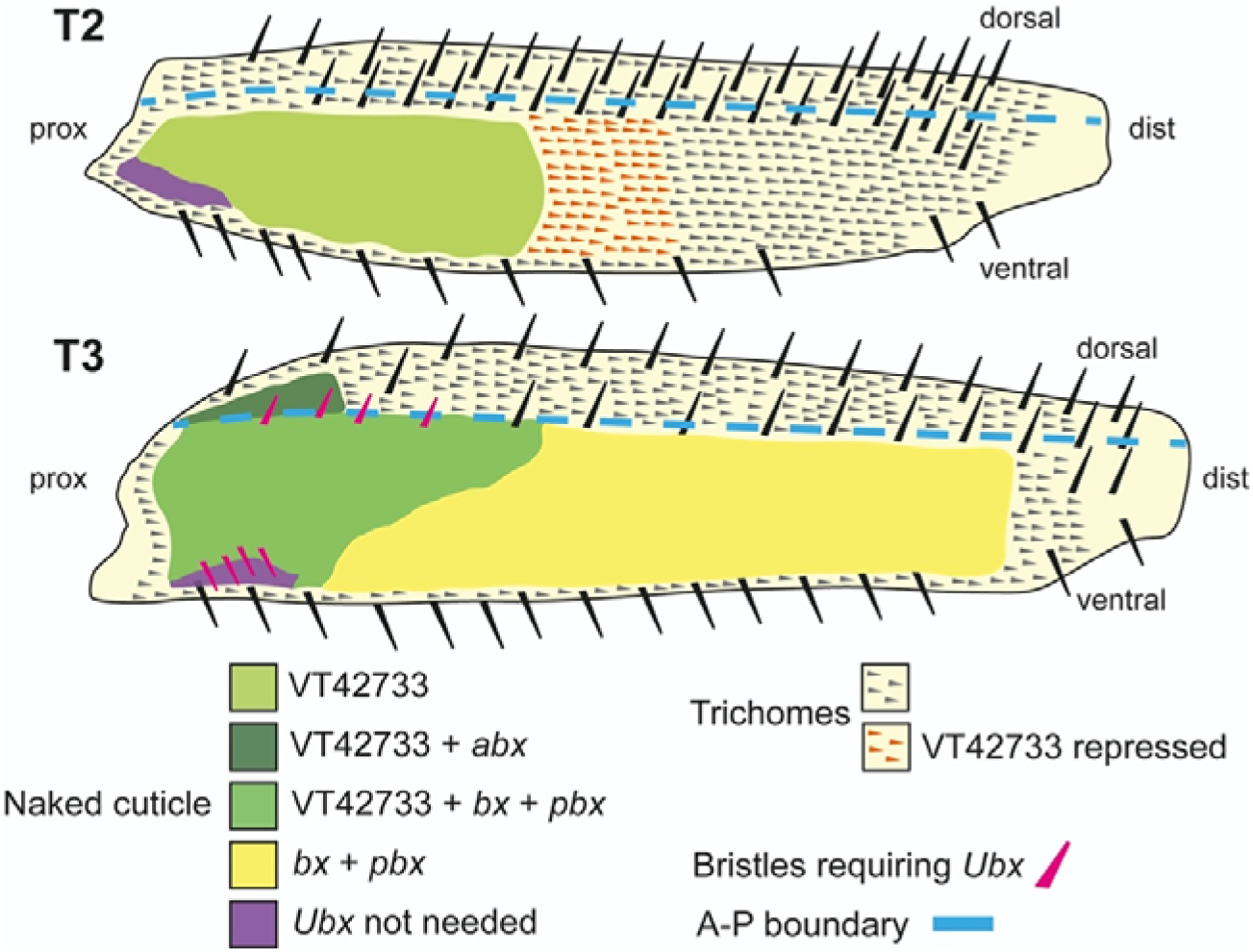
Summary of the regulation and roles of *Ubx* in T2 and T3 femurs. In T2 (upper scheme), *Ubx* expression is regulated in the proximal posterior femur by enhancer VT42733 and, together with the cofactors Exd and Hth, results in the activation of *mir-92a/b* and suppression of trichomes in the so called ‘naked valley’ (light green). Repression of *Ubx* via VT42733 (potentially regulated by *awh, C15, Dll, mirr* or *NK7.1*) more distally in the T2 posterior femur defines the distal limit of *Ubx* expression and the ‘naked valley’ (orange trichomes). In T3 (lower scheme), VT42733 and *abx* regulate *Ubx* expression in the proximal dorsal-anterior of the femur to suppress trichome development (dark green). This also requires *vvl*. In the posterior T3 femur, *Ubx* expression requires VT42733, *bx* and *pbx* proximally (medium green), and *bx* and *pbx* distally (yellow) to suppress trichomes and generate naked cuticle. Patterning of the anterior and posterior T3 femur by *Ubx* also requires *exd* and *hth*. The correct development of small proximal bristles on the T3 posterior femur also requires *Ubx* (pink bristles). In the proximal posterior-ventral femurs of both T2 and T3, trichome development is blocked independently of *Ubx* (purple shading). Scheme based on findings of [23,24,28] and this study.

Davis et al. [23] previously surveyed most of the third intron of *Ubx*, including the VT42733 region, for a leg enhancer (Fig. 2A). However, they did not identify any regions with pupal leg activity although they found that *abx* is required for earlier expression during T2 development consistent with previous studies [23,25,26,38]. This apparent inconsistency with our results could be explained by the different methods used to locate the enhancer. While we used reporter constructs encompassing regions of accessible chromatin in T2 pupal legs to discover that VT42733 is able to drive expression in NV cells, Davis et al. [23] studied this region using deficiencies in trans with *Cbx^3^* and found no effect on the trichome patterning of the T2 femur. This suggests that VT42733 is able to drive expression in femur cells but removal of this region in a trans-heterozygote does not affect the trichome pattern perhaps because of compensation by additional binding sites located elsewhere in the *Ubx* locus. To more directly test this, it would be interesting to precisely delete the leg enhancer from the endogenous location instead of using large deficiencies of the *Ubx* locus that likely have pleiotropic effects and perhaps even result in prepupal lethality. Recent analysis of the *abx* enhancer resulted in similar findings to our study and those of Davis et al. [23]. Delker et al. [20] showed that a reporter construct with a minimal region of 531 bp of the *abx* enhancer is able to recapitulate differential *Ubx* expression in proximal versus distal cells of the developing halteres [20]. However, deletion of this region using CRISPR/Cas9 had no effect on this expression pattern. The authors concluded that there are likely additional binding sites elsewhere and potentially even scattered throughout the *Ubx* locus that contribute to its differential expression in the halteres [20].

It is clear that many of the fragments of the *Ubx* locus that we tested for enhancer activity, including the NV enhancer VT42733, are active in other pupal tissues that express *Ubx* such as the T3 legs but also in places that are not known to normally express *Ubx*, for example the T1 legs. This suggests that these fragments exclude binding sites for TFs that repress *Ubx* in these tissues or other cis-regulatory elements like boundary elements that restrict *Ubx* expression to the correct locations. Ectopic expression has been observed previously with reporter constructs for regulatory regions of *Scr* and *Ubx. abx* fragments drive ectopic expression in imaginal discs that do not normally express *Ubx* and this has been suggested to be a consequence of their exclusion of a nearby polycomb response element [20], and potentially the variably occupied CTCF site in the third intron (Fig. 2A) [36]. Furthermore, reporter constructs for recently identified *Scr* enhancers that reproduce expression of this Hox gene in T1 also appear to be ectopically active in other legs where *Scr* is normally repressed [12]. It was suggested that these reporters contain binding sites that facilitate expression in all legs but they are missing silencer elements that normally restrict *Scr* to T1 [12]. This could also be the case with our *Ubx* leg enhancer. Alternatively, the placement of a cis-regulatory element into a different genomic location could introduce additional TF binding sites or make it accessible in tissues where it is normally inaccessible.

### Candidate TFs for *Ubx* regulation during pupal leg development

Our identification of a T2 enhancer of *Ubx* allowed us to begin to decipher how this Hox gene is regulated in a specific developmental context and to further explore the topology of the surrounding GRN. We tested 25 TFs expressed in pupal legs with predicted binding sites in regions of accessible chromatin in VT42733 and e33.A, and the Hox co-factor Hth (Fig. 4). We found that RNAi knockdown of eight of these factors affected the trichome pattern on T2 femurs (Fig. 4). In contrast Giraud et al. [72] found that only 7/117 TFs they tested had an effect on halteres, which they argued was due to robustness provided by a high dose of Ubx. Our results suggest that the trichome patterning on T2 femurs is more readily genetically perturbed, perhaps because of a lower Ubx dose distally in the femur, which might explain the extensive natural variation in this phenotype [24,61,62].

We found that the known Hox co-factors Hth and Exd are required for Ubx function in T2 and T3 as knockdown of these TFs gave extra trichomes and bristles on dorsal-anterior and posterior femurs similar to *Ubx* loss of function [23,24,28] (Fig. 7). *hth* knockdown had a stronger effect on T2 and T3 than *exd* RNAi, particularly on posterior trichomes and bristles. Indeed, for T3 the effect of *exd* RNAi was restricted to the dorsal-anterior where it resulted in ectopic trichomes. This suggests that Hth is required for Ubx mediated repression of trichomes and bristles in the proximal posterior and proximal dorsal-anterior but Exd is only required in the latter cells (Fig. 7). Given the presence of putative Exd-Ubx dimer binding sites in the VT42733 sequence, this may involve Ubx autoregulation of this enhancer in proximal dorsal anterior cells as shown for Exd-Ubx and Exd-Scr binding in other appendages [20,73,74]. However, Exd-Ubx binding to *abx* represses *Ubx* expression proximally in halteres [20] whereas our results indicate that Exd positively regulates *Ubx* in T2 and T3 femurs.

Apart from Exd and Hth, the only other TF we tested that resulted in an increase in trichomes on T2 and T3 femurs when knocked down was *vvl*. Although *vvl* has no reported role in *Drosophila* leg disc development it is expressed in the growing appendages of other arthropods [75] and our RNA-seq data shows that it is expressed in *Drosophila* pupal legs 24 hAPF during trichome patterning [62]. Our results suggest that *vvl* represses trichomes in the T2 NV and in the dorsal-anterior of T3 perhaps by activating *Ubx* via the putative binding sites in the VT42733 enhancer, although it could act in parallel with or even downstream of this Hox gene.

RNAi knockdown of *awh, C15, Dll, mirr* and *NK7.1* all resulted in an enlargement of the NV on posterior T2 femurs but had no effect on T3. These results suggest that they contribute to repressing *Ubx*, perhaps even directly via their predicted binding sites in the VT42733 enhancer, but again we cannot exclude the possibly that they act in parallel to this Hox gene or downstream.

It has recently been reported that Dll can act as co-factor for Scr to help regulate T1 morphology [73]. Dll and Scr bind to two monomer sites separated by a short space in enhancers of Scr target genes in T1 cells [72]. In T2 and T3 *Dll* is expressed in the coxa and distally in the femurs [76,77]. Dll could therefore also act as a Ubx co-factor to help auto-repress the expression of this Hox gene in T2 and T3 femurs. However, we did not identify any sequences like the Dll-Scr motifs in the VT42733 enhancer suggesting that if Dll does regulate *Ubx* in T2 and T3 it binds as a monomer to some of its eight predicted binding sites in this enhancer to repress *Ubx* expression. It remains possible that Dll acts downstream of Ubx to either activate trichomes distally on T2 femurs or by repressing target genes of this Hox gene that promote formation of naked cuticle. Interestingly, there is evidence that Dll represses other genes during leg development including *serrate* [78].

We suggest that Dll-mediated repression of *Ubx* may help to promote the generation of trichomes on the distal region of the T2 femur while Ubx activates *miR-92a* more proximally to repress trichomes and generate the NV (Fig. 7). However, a more detailed understanding of these regulatory interactions requires assaying whether Dll and Ubx bind directly to the *Ubx* and *mir-92a* enhancers, respectively.

In conclusion, we have identified a leg enhancer of *Ubx* that drives expression to sculpt the fine-scale morphology of T2 and T3 femurs. This provides new insights into the regulation of this Hox gene during postembryonic development and will serve as a platform to better understand how it is wired into the wider leg trichome GRN.

## Acknowledgements

ADB was funded by a BBSRC DTP studentship and SK by a DFG Research Fellowships (Ki 1831/1-1). Stocks obtained from the Bloomington Drosophila Stock Center (NIH P40OD018537) were used in this study. We thank Marianne Yoth for assistance with experiments. This work relied greatly on access to information curated by FlyBase [79].

## Supplementary Figures

**Supplementary Figure 1:**
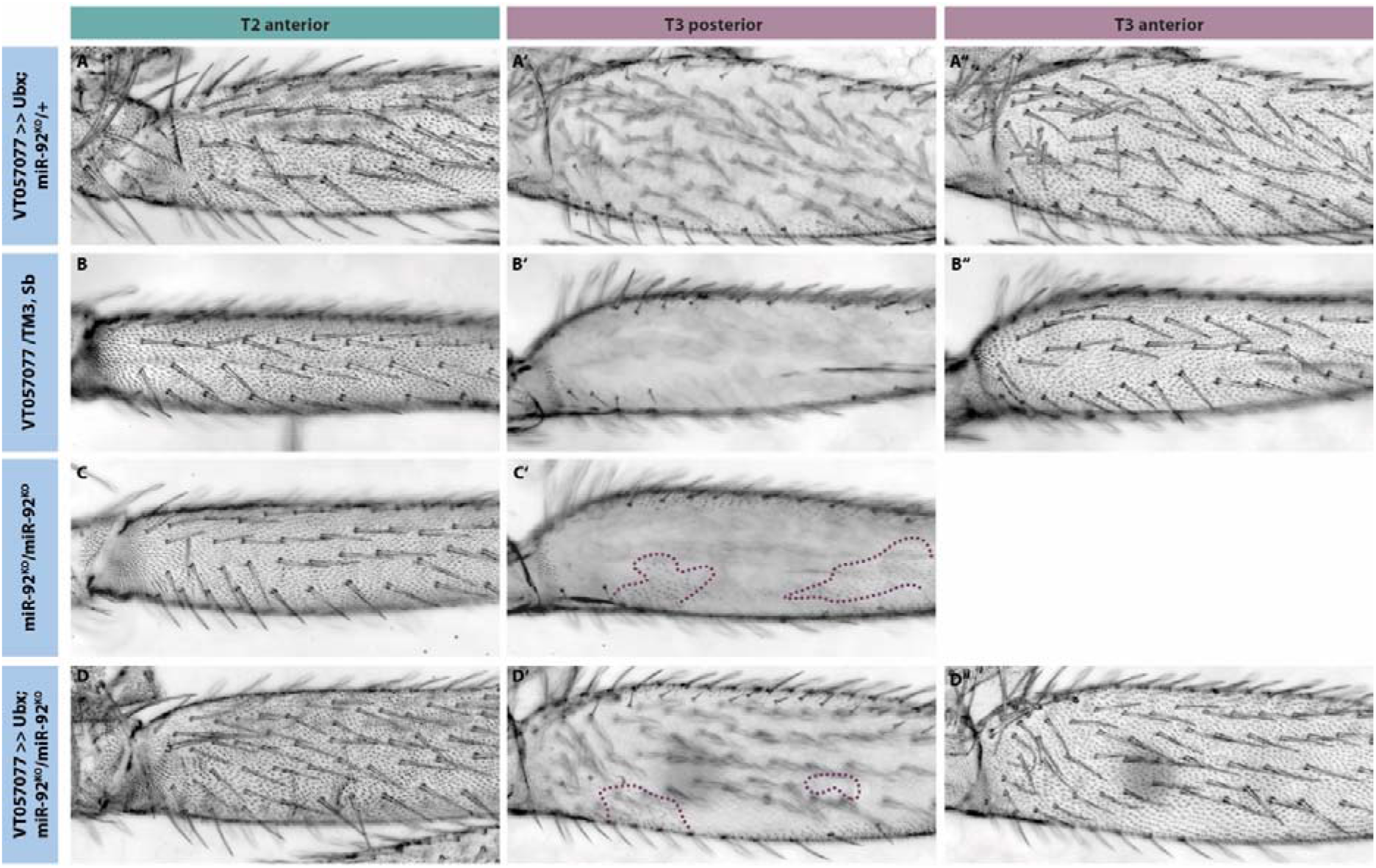
*Ubx* over-expression does not affect the anterior side of the T3 femur and requires *miR-92a* for the repression of ectopic trichomes. (A-A’’’) In contrast to the posterior side of the T2 femur, *Ubx* is unable to repress trichome development on the anterior side of T2 or T3 when overexpressed in the whole leg, and the femurs show no difference to a controls (B-B’’’). (C-C’’) Ectopic patches of trichomes (dashed line) develop on the normally naked posterior side of a T3 femur in a *miR-92* loss-of-function mutant. (D-D’”) These ectopic trichomes are also not repressed by *Ubx* over-expression.

**Supplementary Figure 2.**
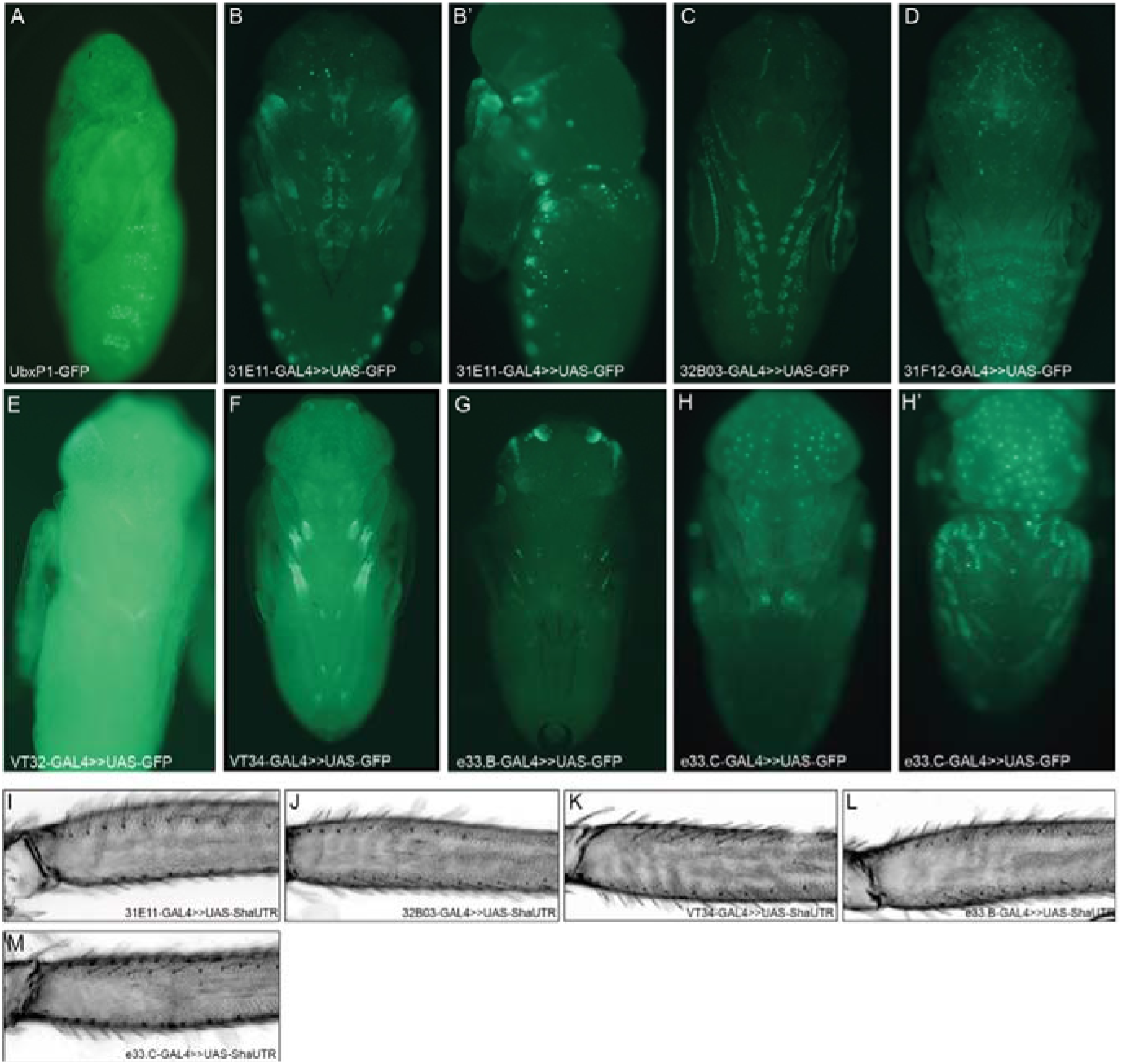
Expression driven by additional reporter constructs in whole pupae at 24 hAPF. (A) Expression driven by the UbxP1 fragment is localised to the abdominal histoblasts. (B-B’) GMR31E11 drives expression in leg joints and in spots along the pupal abdomen. (C) GMR32B03 drives expression along the whole of the leg and also at the periphery of the wing. (D) GMR31F12 drives expression in the abdomen in a stripe like pattern which seems to be located in internal tissues and not the developing epithelium. (E) VT42732 does not drives expression in the pupae. (F) VT42734 drives quite specific expression in all legs. (G) e33.B drives very specific expression in developing legs and also in the pupal antenna. (H-H’) e33.C does drive some leg expression, although this appears to be minimal, as well as expression in the dorsal abdomen. (I-M) The reporter constructs which did drive expression in pupal legs (31E11, 32B03, VT34, e33.B and e33.C) were functionally tested to see if they could induce trichome formation on naked cuticle when crossed to the trichome activating line, UAS-shaΔUTR. However, none of these lines were capable of promoting trichomes in the NV.

**Supplementary Figure 3.**
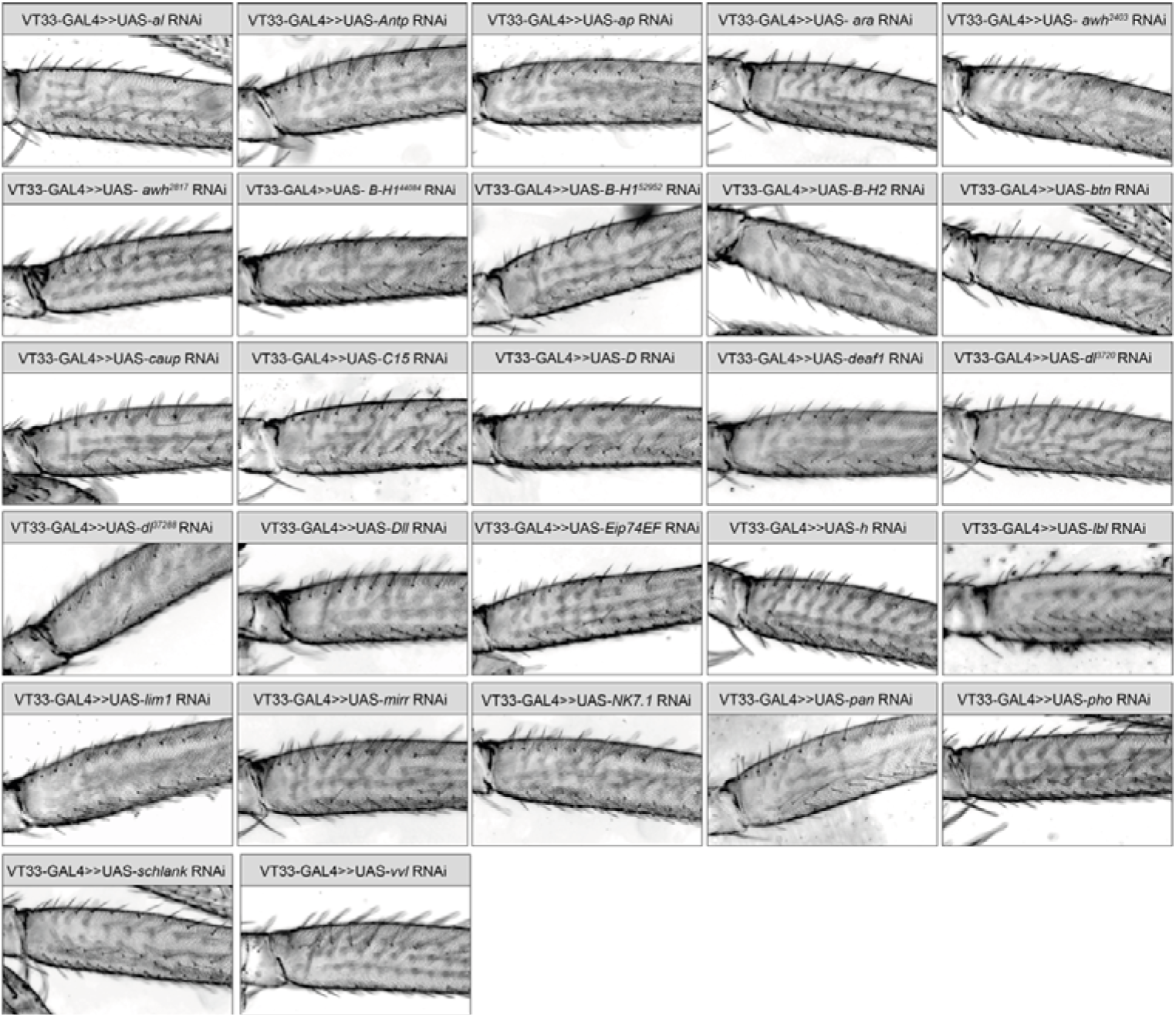
Representative T2 legs from the RNAi screen of predicted TFs. T2 proximal femurs showing the NV for each RNAi line tested.

**Supplementary Figure 4.**
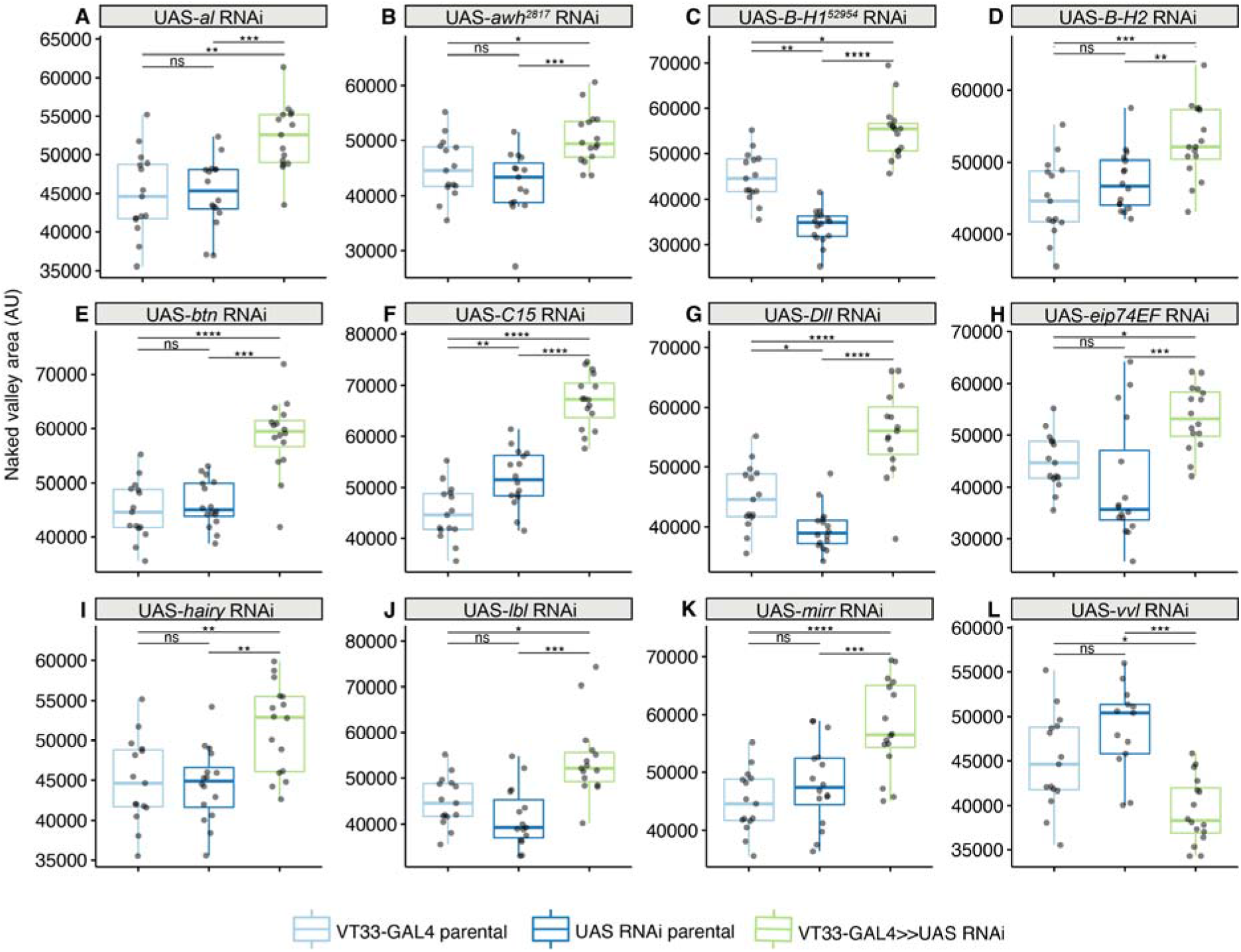
TFs that significantly alter the size of the NV before data normalisation. (A, C, D, E, H, I, J) These TFs significantly increase the size of the NV compared to parental control before normalisation of the measurements. When the data is normalised against the length of the T2 femur these TFs are no longer significant. (B, F, G, K, L) These TFs significantly affect the size of the NV both before and after the measurements are normalised (see Fig.4).

**Supplementary Figure 5.**
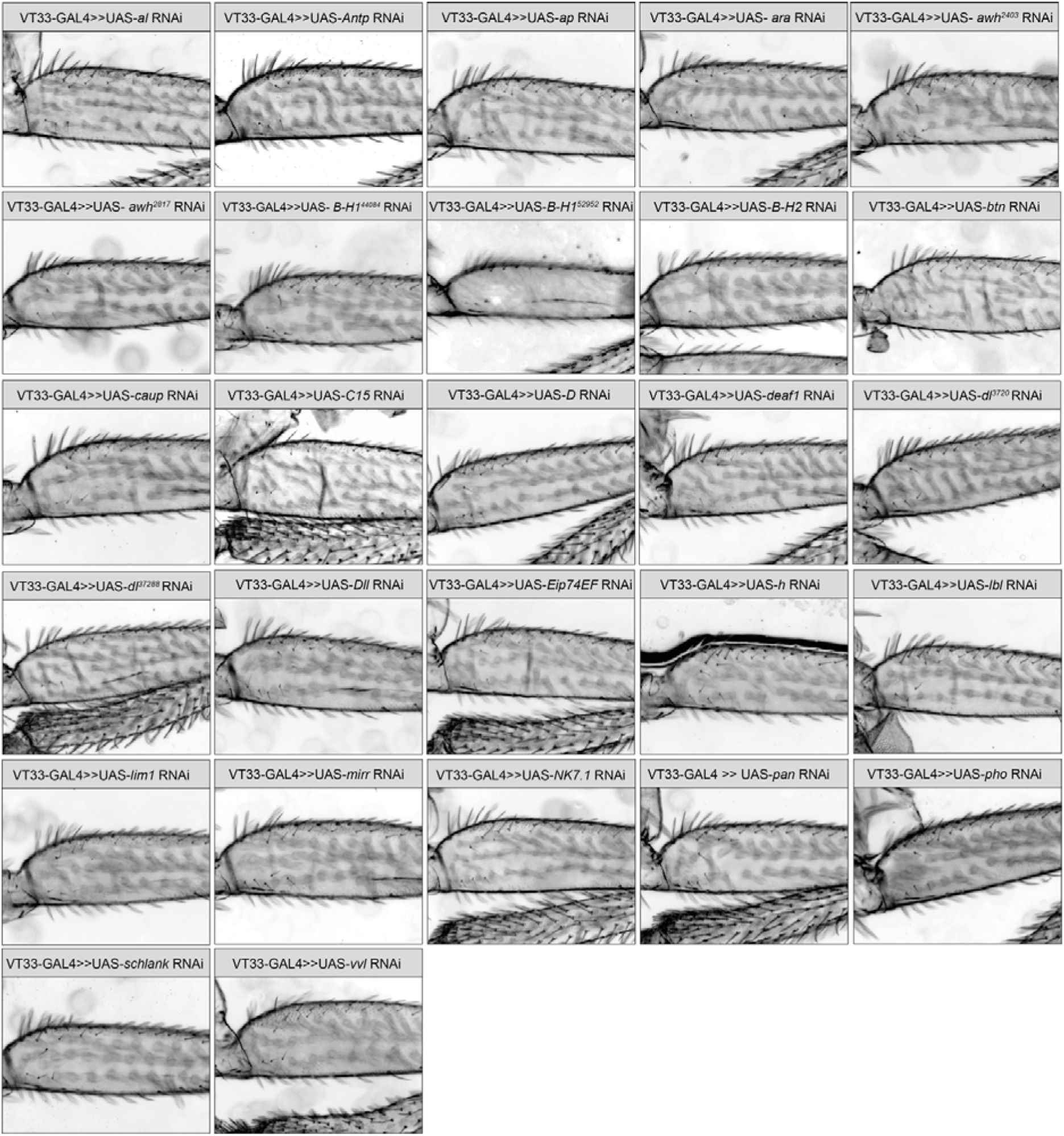
Representative T3 legs from the RNAi screen of predicted TFs. Images of T3 proximal femurs for each RNAi line tested.

Supplementary File 1. A list of all the fly stocks used in this study

Supplementary File 2. Coordinates of reporter lines, primer sequences of reporter constructs and RedFly annotated regulatory regions for the *Ubx* locus.

Supplementary File 3. All TFBS identified by JASPAR including positions relative to open chromatin.

Supplementary File 4. Raw measurements of NV from all RNAi experiments including statistical significance.

